# Adaptive phenotypic divergence in an annual grass differs across biotic contexts

**DOI:** 10.1101/382770

**Authors:** Anna M. O’Brien, Ruairidh J.H. Sawers, Sharon Y. Strauss, Jeffrey Ross-Ibarra

## Abstract

Climate is a powerful force shaping adaptation within species, yet adaptation to climate does not occur in a vacuum: species interactions can filter fitness consequences of genetic variation by altering phenotypic expression of genotypes. We investigated this process using populations of teosinte, a wild annual grass related to maize (*Zea mays* ssp. *mexicana*), sampling plants from ten sites along an elevational gradient as well as rhizosphere biota from three of those sites. We grew half-sibling teosinte families in each biota to test whether trait divergence among teosinte populations reflects adaptation or drift, and whether rhizosphere biota affect expression of diverged traits. We further assayed the influence of rhizosphere biota on contemporary additive genetic variation. We found that adaptation across environment shaped divergence of some traits, particularly flowering time and root biomass. We also observed that different rhizosphere biota shifted expressed values of these traits within teosinte populations and families and altered within-population genetic variance and covariance. In sum, our results imply that changes in trait expression and covariance elicited by rhizosphere communities may have played a historical role in teosinte adaptation to environments and that they are likely to continue to play a role in the response to future selection.

## Introduction

Environmental variation across landscapes is a major selective force driving phenotypic differentiation both within and across species (e.g., Clausen et al., 1947). For example, flowering phenology varies dramatically across latitude in plants from *Arabidopsis* to *Populus* (Stinchcombe et al., 2004; Keller et al., 2012), and climate strongly influences the fitness of species life history phenotypes in common gardens (Rehfeldt et al., 2002; Wilczek et al., 2014). Recent changes in climate have also led to numerous contemporary phenotypic responses, including changes in animal body size (Millien et al., 2006) and plant flowering time (Franks et al., 2007; Willis et al., 2008).

Species interactions are another strong selective force shaping phenotypes. Predator-prey relationships can result in extreme trait escalations (Brodie Jr et al., 2002; Decaestecker et al., 2007; Toju, 2008), and competitive interactions may lead to phenotypic divergence that stabilizes coexistence, such as character displacement or niche partitioning (Pfennig and Pfennig, 2009; Thorpe et al., 2011; Germain et al., 2016). Mutualisms may also alter selection on traits, by either strengthening selection — such as bee pollination causing divergent selection on orchid scent (Ramírez et al., 2011) — or weakening selection, such as loss of function in metabolic traits redundant with functions of mutualistic gut bacteria (Bennett and Moran, 2015).

Biotic and abiotic selective forces may act conditionally: for example, plant-plant interactions often shift from negative to positive under increasingly stressful abiotic conditions (Callaway et al., 2002), and plants may exclude fungal root symbionts in high nutrient soils (Treseder, 2004; Grman, 2012), or the degree of evolutionary trait escalation may depend on environment (Toju et al., 2011; Stokes et al., 2015; O’Brien et al., 2018). Interactions can even be gained or lost with changes in climate, such as expulsion or death of endosymbionts at high temperature in insects (Wernegreen, 2012) and corals (Hoegh-Guldberg, 1999), or through phenological mismatches, such as in plant-pollinator interactions (Burkle et al., 2013). In short, the interdependence of abiotic and biotic selection on trait differentiation may be pervasive.

Not all changes to phenotypes are caused by selection; plasticity in trait expression in response to changes in the biotic or abiotic environment is ubiquitous (e.g. Falconer, 1952; West-Eberhard, 1989), and linked to widespread differential effects of genotypes across environment (Hunter, 2005; Des Marais et al., 2013). A key set of biotic environments that lead to plasticity in plant traits are rhizosphere biota, which alter the expression of a wide range of phenotypes (Friesen et al., 2011; Goh et al., 2013). Rhizosphere biota are the collection of bacteria, nematodes and fungi living in the vicinity of plant roots (the rhizosphere, Hiltner, 1904; Bais et al., 2006). Species composition in these communities is influenced most strongly by abiotic factors, but plant genotype also contributes (Bulgarelli et al., 2012; Peiffer et al., 2013; Bouffaud et al., 2014; Lebeis et al., 2015; Walters et al., 2018).

Biota-mediated trait expression may play a critical role in adaptation as plant populations encounter new environmental conditions. Expression of plant traits is known to depend on plant genotype, biota species composition, the environment in which these interactions take place, and interactions among these properties (Klironomos, 2002; Zhu et al., 2009; Johnson et al., 2010; Smith and Read, 2010; Lau and Lennon, 2012; Wagner et al., 2014; Rúa et al., 2016). Indeed, rhizosphere biota are already implicated in current range shifts (Lankau et al., 2015), in species invasions (Hayward et al., 2015), and in trait responses to experimental selection on plants and soil biota for plant drought tolerance (Lau and Lennon, 2012) or flowering time (Panke-Buisse et al., 2015).

Here, we test the importance of plant-biota interactions in the expression of adaptive phenotypic divergence and genetic variation using interactions between a subspecies of teosinte and its rhizosphere biota. The common name teosinte includes several subspecies of wild annual grass related to domesticated maize (*Zea mays* ssp. *mays*) that are distributed throughout Mesoamerica (Hufford et al., 2012). We focus our analysis on the highland teosinte subspecies *Zea mays* ssp. *mexicana*, a close relative of maize (divergence ≈ 60,000 years ago, Ross-Ibarra et al., 2009) which still hybridizes with maize where they co-occur (Hufford et al., 2013), and when grown in the same soil, hosts similar rhizosphere biota (Bouffaud et al., 2014; Szoboszlay et al., 2015). In fact, genetic divergence between maize and teosinte subspecies also predicts divergence in their rhizosphere biota communities (Bouffaud et al., 2014) and adaptation to local rhizosphere biota has been documented in teosinte populations (The authors, 2018) over smaller timescales. Teosinte exhibits a number of phenotypes that are known to differ along elevational gradients and are suspected to be adaptive, including phenology (Eagles and Lothrop, 1994; López et al., 2011), plant architecture, plant size, and stem color (Doebley, 1984; Lauter, 2004; Hufford et al., 2013). To assess the role of rhizosphere biota in the expression of adaptive teosinte phenotypes to climate, we ask whether 1) interactions with rhizosphere biota alter how phenotypes are expressed, 2) whether teosinte shows evidence of adaptive phenotypic divergence patterned by climate in rhizosphere-altered traits, and 3) whether rhizosphere biota alter the potential future evolutionary responses of teosinte.

## Methods

### Plant and biota sources

We used seed and biota collected from 10 teosinte populations from central Mexico in 2013. Information on these populations indicates differences among sites in both climatic conditions (Bioclim, Hijmans et al., 2005, extracted with package raster Hijmans, 2015 in R, R Core Team, 2014) and soil characteristics (see The authors, 2018). The sites ranged 6.6 °C in mean annual temperature (MAT), more than 1100 meters in elevation, from sandy to clay soil, and the wettest site received nearly twice the annual precipitation of the driest site (The authors, 2018). We randomly selected three of these sites to use as sources of rhizosphere inocula: San Mateo Tezoquipan, San Matías Cuijingo, and South Toluca. We refer to biota sources throughout using the mean annual temperature at the site (they become Biota15.0, Biota14.3, and Biota13.0, respectively). Biota15.0 and Biota14.3 are separated by 15.4 km, Biota15.0 and Biota13.0 by 96.1 km, and Biota14.3 and Biota13.0 by 94.6 km.

In August 2013, teosinte rhizosphere soil and roots were collected from the rhizosphere of adult plants at each site by unearthing roots, shaking off loose soil, collecting the remaining soil and roots, and repeating with dispersed plants from the site (≈15) until the material totaled 2 kg. Rhizosphere soil and roots were kept refrigerated at 4°C until used in the experiment, when samples were homogenized in a blender. These collection and storage procedures were designed to maintain viability of both bacteria and fungi.

### Experiment

In July 2014 we planted seeds from each population, inoculating them with one of the three rhizosphere biota sources. For each combination of plant population and biota source, we planted 3 pots with seeds sampled from separate inflorescences from each of 10 mature plants (30 total pots per population × biota combination). Different female inflorescences likely sample pollen from different pools of possible fathers for a few reasons: selfing rates in teosinte are likely low (Hufford et al., 2011, ≈ 3% in the related ssp. *parviglumis*) and we observed in the field both that inflorescences on the same plant matured at different times and that each plant had only one male inflorescence (first author, personal obs). We therefore treat the 3 seeds from each maternal plant as half-siblings.

We grew plants in 2.83 L pots (Stuewe & Sons Treepots) with steam sterilized (4 hours at 93°C using a PRO-GROW SS60) potting mix (90% sand, 5% perlite 5% vermiculite 0.2% clay). To inoculate, we filled pots to 2 L with sterilized mix, added 50 mL of a 4:1 homogenized mix of sterile sand and inocula, and filled to the top with sterilized mix. We added seeds to pre-watered pots after scarification and overnight soaking. We randomized the bench planting design with respect to seed source, inoculum source, and maternal family. We added up to three seeds to a pot as supplies allowed, recorded the date of germination for all seeds, and weeded after germination if more than one plant germinated.

To encourage germination, we kept pots moist and unfertilized for two weeks, then watered and fertilized once per week with Hoagland’s low P. As plants grew and demands of plant tissue for water increased, we increased water from 100 mL per week to 200 mL per week for the last 4 weeks. However, the total amount of fertilizer applied to each plant was constant, such that we applied phosphate ion at a rate of 100 *µ*mol per week (at first in 50 *µ*M solution then decreasing to 25 *µ*M).

Plants began flowering in September, and we recorded first flowering date when silks or anthers were first visible. We harvested each adult plant 15 days after its first inflorescence was observed. At harvest, several additional phenotypes were measured: stem width at the highest node from which aerial roots contacted the soil, the height from soil to highest ligule, the width of the penultimate leaf subtending the primary male inflorescence (tassel), and the length of the primary axis of the tassel. A photograph of the stem was taken with a color standard, from which greenness of the oldest pre-senescence leaf sheath was measured using ImageJ (Schneider et al., 2012) and corrected as suggested in (Stevens et al., 2007). Plant roots and shoots were separated at the highest node where roots entered the soil, dried at ambient temperature until mass stabilized, and weighed. For each of these traits, we expected variation might be of adaptive importance to teosinte due to previous speculation in the literature (Doebley, 1984; Eagles and Lothrop, 1994; Lauter, 2004; López et al., 2011; Hufford et al., 2013) and obvious differences across field populations (the authors, personal obs, Figure S1).

### Genotyping

An additional 9 seeds from each population were grown in a greenhouse at the University of California Davis. Young leaf tissue was sampled for DNA extraction using the DNeasy Plant Mini Kit from Qiagen. A single-nucleotide polymorphism dataset was generated from genotype-by-sequencing (GBS) (Elshire et al., 2011; Glaubitz et al., 2014) at the Biotechnology Resource Center, Cornell University, generating low coverage data at 955,690 SNPs. The GBS dataset was filtered to sites with data across at least 86 of the 90 plants (95% coverage), resulting in 60,377 SNPs. We removed two individuals with more than 70% missing data (remaining individuals averaged 0.9 % missing data, ranging from 0.2% to 4.2%).

### Effects of biota on trait expression

We first asked whether biota affect the expression of our set of putatively adaptive teosinte phenotypes. Using experimental trait data we fit and compared linear model structures for main effects and random effects. We tested four random effect structures: 1) rhizosphere biota treatment, family, and population, all separately, 2) additionally including population within biota treatment, 3) additionally including family within biota treatment, or 4) additionally including both family and population within biota treatment. We also fit models with main effects, either: 1) intercept only or 2) intercept and a main effect of one environmental variable for the plant population source. For this environmental variable, we compared elevation (ELV, masl), mean annual temperature (MAT, °C), total annual precipitation (TAP, in millimeters) or soil water holding capacity (SWC, weight percentage of water held at saturation at 0.3 bars). Each trait and explanatory variable was centered on the mean and scaled by the standard deviation. We ran models with MCMCglmm (10 chains of 100,000, burn-in 20,000, thinning 100, Hadfield, 2010) in R (R Core Team, 2014) and compared models using the deviance information criterion (DIC, Spiegelhalter et al., 2002).

### Divergence of trait means across teosinte populations and environments

We tested whether teosinte traits have adaptively diverged both across populations in general and specifically in response to environmental variation. To develop a neutral expectation for trait divergence, we estimated coancestry (expected relatedness of a pair of individuals) both within each population and between all pairs of populations using our SNP dataset. We computed coancestry between populations (Karhunen and Ovaskainen, 2012, using the package RAFM in R) with a random subset of 10,000 loci from the GBS dataset and parameters recommended by the package authors (20,000 iterations, 10,000 burnin, and thinning by 10).

Pairing coancestry estimates with ancestral trait variance-covariance estimates allowed us to quantify the expected magnitude of trait shifts during population divergence due to neutral processes alone. We estimated ancestral trait variance and covariance for all 9 traits using Driftsel (Ovaskainen et al., 2011; Karhunen et al., 2013) in R (R Core Team, 2014). Driftsel leverages phenotype information in related individuals and pairwise population coancestry to generate neutral expectations for trait mean shifts in the full set of traits. It then uses differences from these expectations across populations and traits to evaluate the effect of selection on the divergence of trait means across populations. Both here, and for all further trait analyses, we mean-centered phenotype data and scaled by the standard deviation, as recommended by Hansen and Houle (2008). We ran Dritfsel for 440,000 iterations with a 40,000 iteration burn-in and thinning by 2,000 (determined by increasing iterations until MCMC samples converged). We used weak priors recommended by the Driftsel authors (Karhunen et al., 2013). We performed the test of trait divergence for data from each inoculum treatment separately. We first focused on results of this test for all traits collectively (S statistic), which accounts for predicted co-drift of trait means due to the structure of the ancestral trait variance and covariance (G) matrix. The S statistic ranges from 0 to 1: values below 0.05 indicate strong evidence for stabilizing selection, values near 0.5 imply perfect drift, and values above 0.95 indicate strong evidence for divergent selection (Ovaskainen et al., 2011; Karhunen et al., 2014). Since only some traits may be under divergent selection, we then evaluated individual traits using only ancestral means and additive genetic variance. We also briefly explored divergence in bivariate trait space to illustrate how considering multivariate space alters expectations. We compared the results across inocula treatments to assess the contribution of effects of rhizosphere biota on trait expression.

We then tested whether the pattern of trait divergence among populations was correlated with abiotic conditions, above and beyond an expected correlation owing to the genetic relatedness among populations. Such a result we would interpret as support for local adaptation to environmental conditions driving phenotypic shifts. We combined the results from Driftsel and environmental data to perform the H test (Karhunen et al., 2014), which pools information across traits, environmental variables, and genetic variation. We did two sets of analyses: first, a set of three key environmental variables (MAT, TAP, and SWC), and second, combining the first two principal components of the Bioclim variables and first two principal components of the soil variables reported in The authors (2018), which each include many co-correlated measures and may be a comprehensive summary of environmental variation. An H greater than 0.95 indicates significant correlation of phenotypes and environment beyond what population genetic similarity alone would predict (Karhunen et al., 2014). We performed H tests for habitat driven trait divergence in each biota treatment separately.

At low elevation, hybrids between highland teosinte and the related low elevation subspecies (parviglumis, *Zea mays* ssp. *parviglumis*) can form (Pyhäjärvi et al., 2013), whereas at higher elevation, introgression from maize (*Zea mays* ssp. *mays*) can be common (Hufford et al., 2013). Introgression could neutrally increase both phenotypic and genotypic divergence, or could be a source of genetic material underlying adaptive divergence. We evaluated population structure between our 10 populations and known individuals of ssp. *mays*, ssp. *mexicana*, and ssp. *parviglumis* to see if gene flow across subspecies could be a source of adaptive genetic material. Additional GBS data from Swarts et al. (2017) were filtered for overlapping SNPs with our dataset (41,385 SNPs), and subset to include approximately equal numbers of individuals from each subspecies (153, 141, 144, of *mays*, *mexicana*, *parviglumis*), using VCFtools (Danecek et al., 2011), PLINK (Purcell et al., 2007), R (R Core Team, 2014), and Ensembl Assembly Converter (Kersey et al., 2017). We then analyzed this dataset with PCA in TASSEL (Bradbury et al., 2007; Glaubitz et al., 2014) to assess genetic relationships.

### Variation in G matrices and response to selection across teosinte population and rhizosphere biota

To test whether interactions with rhizosphere biota might influence future responses to selection, we estimated G matrices representing the additive genetic variance (diagonal elements) and covariance (off-diagonal elements) of traits. We performed G matrix estimation for each population in each rhizosphere inoculation treatment separately (data from 30 plants), for the 5 traits that showed evidence of divergence across populations. We fit animal models on centered and scaled trait data (as above). Briefly, animal models assume individual phenotypes (vector *y_i_*) are functions of the mean trait value (*µ*), additive genetic breeding values (*a_i_*) and residual effects (*e_i_*): *y_i_ ∼ µ*+*a_i_* +*e_i_*. Linear model fitting of individual phenotypes to the animal model further assumes breeding values fit the genetic variance-covariance matrix *G*, where elements of *G* rest on the half-sibling covariance (e.g. diagonal genetic variance *V_A_* elements are four times the estimated covariance of that trait among half-siblings, see Falconer and Mackay, 1996). Together with a residual error variance-covariance matrix *R*, *P* the phenotypic variance-covariance matrix among traits will then be: *P ∼ G* + *R* (see more thorough and applied explanations in Lynch et al., 1998; Wilson et al., 2010).

We fit models with MCMCglmm() in R (Hadfield, 2010; R Core Team, 2014). We assumed normally distributed traits, used random effects for trait means, and applied a weakly informative inverse Wishart prior, biased towards very low additive genetic variance and covariance (Hadfield, 2012). We fit models for 1,000,000 iterations, with 100,000 burn-in and thinning by 100. We checked MCMC chains for convergence (traces, see Figure S7) and implemented two alternate priors: reduced expected variance or both reduced expected variance and weaker bias (Hadfield, 2012). All priors yielded visually similar G matrices and similar results for subsequent analyses, so we only present results for the recommended prior.

To test the significance of variation we observed in the G matrices across populations and rhizosphere inoculation treatments, we used eigentensor analysis (Aguirre et al., 2014). This analysis first calculates the variation among the set of 30 G matrices for each cell in the matrix and covariation for each pair of cells, and collects them into a summary variance-covariance matrix. For sets of G matrices where each G matrix has *n* distinct values (ours have 15, see Figure 6), this summary variance-covariance matrix includes *n × n* cells, for covariances between each pair of cells in the G matrices, with *n* cells for variances within cells in the G matrices. Eigentensor analysis then uses eigendecomposition (used also in principle components analyses, for example) of this variance-covariance matrix to find the major axes of variation among the G matrices, known as 4th order tensors, or eigentensors. Each eigentensor contains the contribution of each cell of the G matrix to that eigentensor, e.g. to the variation among the G matrix set explained by that eigentensor. We further decomposed the eigentensors into eigenvectors, which quantifies the influence of individual traits (across all cells of the G matrix in which they are found) on the eigentensor. All eigentensor analyses include uncertainty in the estimation of the real G matrices by including the full posterior MCMC estimates. We performed the analysis in R using code from Aguirre et al. (2014) and the packages matrixcalc and gdata (Novomestky, 2012; Warnes et al., 2014).

To test significance, we compare the amount of variation among the G matrices that is explained by eigentensors estimated on the real and randomized sets of MCMC estimates. Using the breeding values estimated by the animal models, we randomized individual MCMC estimates of breeding values across the population and pedigree information and used each MCMC randomized breeding value set to generate MCMC samples of randomized sets of G matrices, and re-ran the eigentensor analysis across the MCMC samples. If eigentensors calculated from real data explain more variation than eigentensors calculated from randomized breeding values, this indicates significant variation in the real set of G matrices (Aguirre et al., 2014).

We used a selection gradient approach to predict what responses teosinte traits would have to a certain selection gradient (abbreviated *β*). *β* is a vector representing selection on each of the traits in the G matrix. Our *β* was for strong selection on days to flowering (days to flowering perfectly correlated to relative fitness, and selection of one standard deviation change in mean phenotype before reproduction), which we chose due to its high influence on the first eigentensor (see Results). We then calculated the predicted response 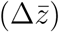 of another trait with high influence on the first eigentensor (see Results) across the posterior distribution for each G matrix, using the multivariate breeder’s equation: 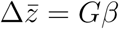 (Lande, 1979). We visualized responses for each estimated G matrix, and assessed overlap with bivariate density kernels using the package ks (Duong, 2018) in R (R Core Team, 2014). We scaled and centered traits prior to fitting G matrices, so responses are standard deviation units relative to a mean at 0.

## Results

We aimed to examine the role of biotic interactions in phenotypic divergence across abiotic environments in the highland teosinte *Zea mays* ssp. *mays*. We grew a common garden experiment of 10 teosinte populations in three separate rhizosphere biota treatments and measured a set of putatively adaptive phenotypic traits, including phenology, vegetative morphology, and color (Doebley, 1984; Eagles and Lothrop, 1994; Lauter, 2004; López et al., 2011; Hufford et al., 2013). Using environmental and genotype data from the same populations, our approach tested whether biota treatments affect trait expression, whether trait divergence is shaped by adaptation to environmental gradients, and whether biota alter expressed genetic variation.

### Rhizosphere biota alter the expression of adaptive phenotypic variation

Visual inspection of all phenotypes shows variable reaction norms across populations and families to different biota (Figure 1, slope steepnesses and signs) and variable means across populations (Figure 1, height of lines). Using the full set of trait data, we fit linear models for each phenotype with a range of random effects testing for genotype-by-biotic environment effects as well as fixed effects including intercept only or source site abiotic environment (see Methods). While there is often no difference in fit between the best model and the next best model for most traits (Table S1), we can nonetheless make some generalizations.

**Figure 1:**
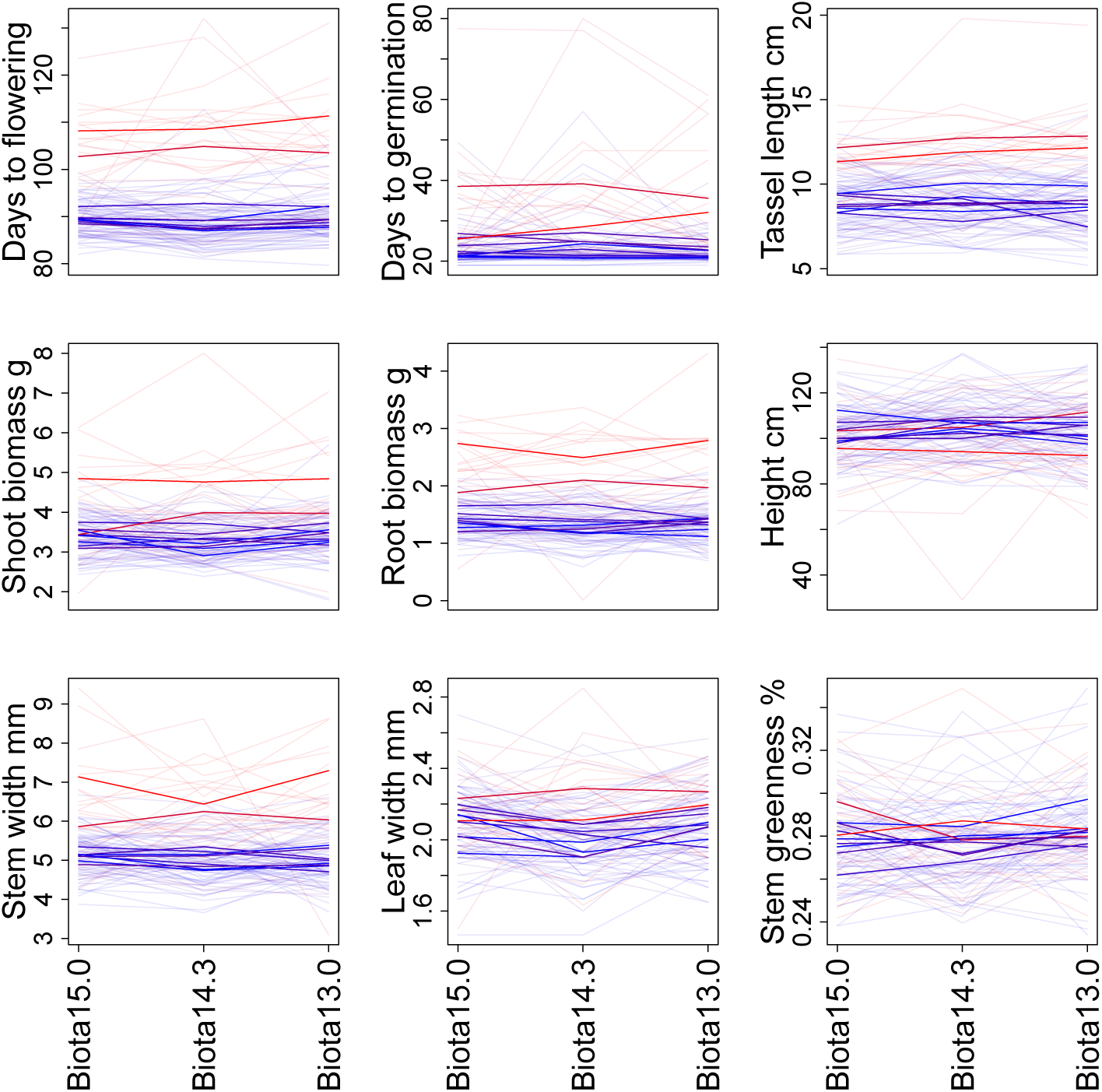
Average trait values (y-axis) for each trait (plots) in each biota (x-axis). Populations (bold lines) and families (faint lines) are both presented and are colored according to mean annual temperature at their source site. Redder indicates populations from warmer sites; bluer indicates populations from colder sites.

Some putatively adaptive teosinte traits we selected for study not only varied plastically across biota, but plastic responses depended on the family or population of plants. Models with genotype-by-biota (for population, family, or both) interactions outperformed models without when ranked by deviance information criterion (DIC) for several traits (days to germination and to flowering, stem and leaf width, and root biomass, Table 1, Figure 1).

**Table 1:** Best models for traits, including best environmental variable at plant source site and random effect structure based on DIC comparisons. When no environmental variable (MAT, mean annual temperature; SWC, soil water holding capacity) is in the best model, cells are left blank (annual precipitation and elevation were never in the best model). F indicates family-by-biota random effect, P is population-by-biota random effect. DIC base indicates the simplest model (intercept and separate random effects for rhizosphere biota, family and population, but no GxE effects). **: pMCMC < 0.05,*: pMCMC < 0.1.

| Trait | Main | Random | Slope | DIC | DIC base |
| --- | --- | --- | --- | --- | --- |
| Days to flowering | – | F | – | 1713.0 | 1713.4 |
| Days to germination | SWC | F + P | 5.1e-3** | 2131.6 | 2157.6 |
| Tassel length | SWC | – | 8.5e-3** | 2096.6 | 2096.8 |
| Shoot biomass | – | – | – | – | 2233.3 |
| Root biomass | – | F + P | – | 1888.7 | 1889.6 |
| Height | MAT | – | 0.11 | 2344.1 | 2444.7 |
| Stem width | – | F + P | – | 2083.4 | 2083.9 |
| Leaf width | MAT | P | 3.3e-3** | 2309.1 | 2310.0 |
| Stem greenness % | – | – | – | – | 2261.6 |

Furthermore, some trait means varied across population source environment, validating expected patterns of trait correlations. The best models for several of the traits included significant slopes with environmental variables (pMCMC < 0.05, days to germination, tassel length, leaf width, Table 1). Though the best models for other traits (stem greenness, stem width, root biomass, shoot biomass, and days to flowering) lacked slope terms with environment variables of the source site, terms were sometimes significant in models with higher DIC (Table S1). Mean annual temperature and soil water holding capacity were the explanatory environmental variables included in the best models explaining trait differences among populations (Table 1). Differences in DIC across variables are slight, however (Table S1), suggesting equally strong links between these site descriptors and population trait differences.

### Trait divergence in teosinte populations is the result of adaptation to environment

To build our null hypothesis for levels of trait divergence, we first estimated genetic drift with coancestry, the probability that alleles chosen from two individuals are identical by descent. Our estimates indicated that most populations were equivalently related (coancestry between populations mostly ≈0.1) and have little drift or inbreeding (within population coancestry only slightly higher, Figure 2). However, two populations were not genetically similar to any other population (coancestry < 0.05), and had higher within population coancestry (≈0.2), indicating more inbreeding and drift away from all other populations.

**Figure 2:**
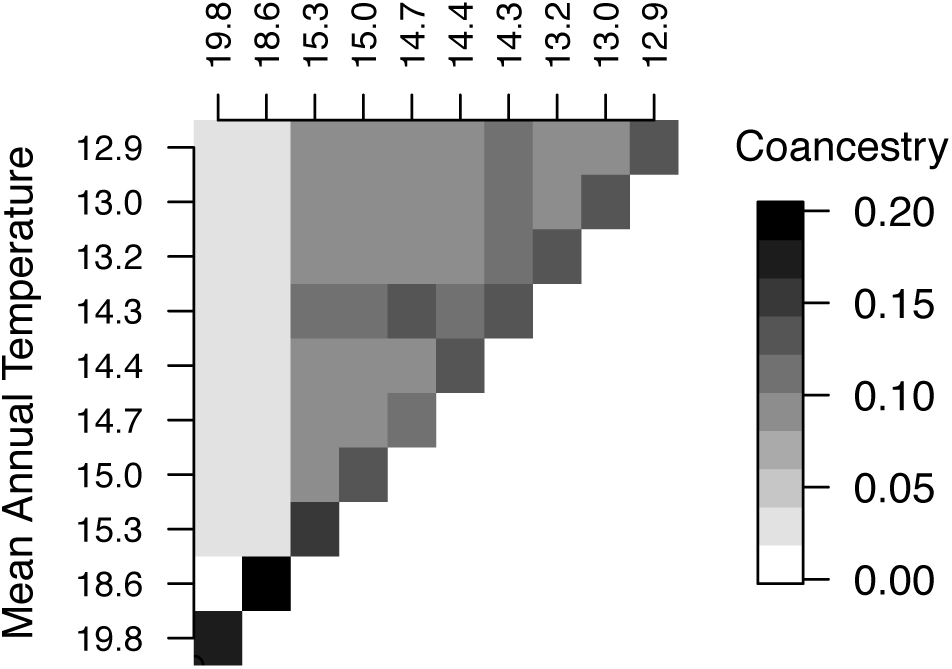
Coancestry within and between all teosinte populations, identified here by the mean annual temperature at their source site.

We tested whether the divergence of trait means across teosinte populations reflects neutral drift or selection using Driftsel (Ovaskainen et al., 2011; Karhunen et al., 2013) to compare divergence in phenotype to expectations based on our observed patterns of genetic coancestry. We performed these comparisons separately for each rhizosphere biota in order to assess the contribution of rhizosphere interactions to our ability to detect the signature of adaptation.

We found that a subset of putatively adaptive traits have diverged more than expected based on genetic coancestry. The statistic S ranges between 0 and 1, and summarizes divergence across the G matrix as a whole (i.e. across all traits simultaneously), where values closer to 1 indicate support for divergence. The biota produced S statistics 0.73, 0.87, and 0.77, suggesting only weak support for divergence of populations across the full trait dimensionality in excess of neutral drift from the ancestral means and G matrix (S greater than 0.95 would indicate strong support, Ovaskainen et al., 2011; Karhunen et al., 2014). However, relative to the expectations from ancestral means, additive genetic variance and drift, two traits (days to flowering and root biomass) individually exceeded expected divergence in at least one population in all three biota sources. Three more traits (days to germination, shoot biomass, stem width) individually exceeded expected divergence in at least one population in only one or two biota sources (Figure 3). Four traits (tassel length, plant height, leaf width, and stem greenness) never exceeded drift expectations for the univariate case (Figure 3). Populations varied in divergence as well, with only two populations exceeding expectations for one or more traits across all biota, and three populations falling within expectations of phenotypic drift for all traits in the univariate case (Figure 3). More traits and population combinations fell beyond neutral expectations when compared in two or more dimensions (see Figures S2, and S3).

**Figure 3:**
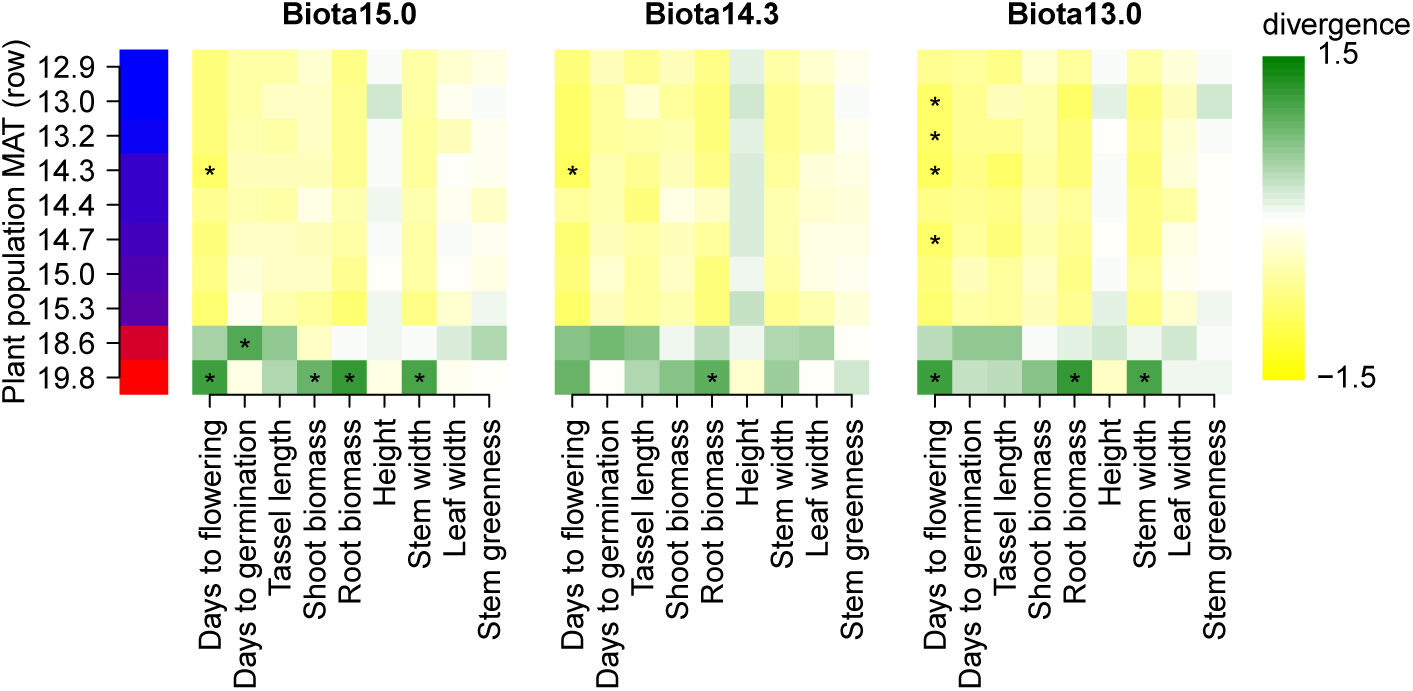
Population divergence from the ancestral mean breeding value for all populations (populations in rows, sorted by MAT at source site) in each biota (separate plots). Green indicates lower values for traits than the ancestral mean breeding value, while yellow indicates higher values. Populations and traits that exceed the 95% confidence interval for neutral divergence are marked with asterisks. More populations and traits exceed expectations in multidimensional space (see Figure S2 and Results).

In contrast, when we considered the pattern of trait divergence across habitat, we found strong evidence that environment shaped trait divergence across populations. If habitat similarity explains significant trait similarity among populations after accounting for genetic similarity of populations in similar habitats, this provides evidence for non-neutral trait divergence across habitat (Karhunen et al., 2014). The “H” test in Driftsel (Ovaskainen et al., 2011; Karhunen et al., 2013) compares the similarity of the population coancestry, habitat similarity, and phenotype similarity matrices, and asks whether the habitat similarity matrix explains variation in the phenotype similarity matrix after accounting for population coancestry. Specifically, if teosinte populations from similar habitats are more similar phenotypically in the common gardens than would be expected from their genetic similarity, this is evidence that environmental variables have shaped trait divergence. The H statistic ranges from 0 to 1, with values over 0.9 or 0.95 indicating 90% and 95% confidence that environment selected on phenotypes, respectively. We observed highly significant H statistic values of 0.995, 0.985, and ≈1 in each biota for our selected environmental variables (for the more environmentally wholistic principal component variables, H statistic values are equally significant: ≈1, 0.995, and 0.980). This result can be intuited by inspecting the estimated population effects (Figure 3) when compared to coancestry in Figure 2: despite the fact that the two low elevation, warmer-sourced populations are genetically very different from each other, they have very similar trait values for most traits, and these trait values are opposite those of all higher elevation, colder-sourced populations. In follow-up analysis without these two populations, we did not detect a signal of phenotypic divergence patterned by climate in any biota (highly non-significant S statistics 0.42, 0.43, 0.38, and H statistics 0.42, 0.30, 0.30, for Biota15.0, Biota14.3, and Biota13.0, respectively).

One possible explanation for these results is that hybridization with one or more of the other two local subspecies of *Zea mays* could have shaped both coancestry and trait divergence. To test this hypothesis, we compared our genotypic data to public SNP datasets for three different subspecies of *Zea mays*. Principal component analysis of SNP genotypes produced three clusters corresponding to the three subspecies (Figure 4), and also showed that both lowest elevation populations from our study have slightly increased genetic similarity with ssp. *parviglumis*, the lowland subspecies, relative to our other populations. Specifically, our low elevation populations have slightly more similarity to ssp. *parviglumis* on the second PCA axis and to particular *parviglumis* populations on the fourth axis (Figure 4, top two plots). On axis 1, however, they remain clearly united with the other highland teosinte (ssp. *mexicana*) populations (4, lower plots). Together with our above coancestry analysis, which indicates that our lower elevation populations have low coancestry with each other (Figure 2), the PCA results suggest that if these populations have gene flow from ssp. *parviglumis*, it is with different ssp. *parviglumis* populations. For some traits, our low elevation population have means shifted in the direction of ssp. *parviglumis*, which has greener stems (Doebley, 1984), is shorter, has later flowering (Sánchez González et al., 1998), later germination (López et al., 2011), and probably greater root mass (Gaudin et al., 2014) than ssp. *mexicana*, but in contrast has shorter male inflorescences (Iltis and Doebley, 1980), and narrower leaves (Sánchez González et al., 1998). In sum, while our low elevation teosinte populations do not appear to be extensively hybridized with the lowland subspecies, they may have independent rare or old hybridization events with different ssp. *parviglumis* populations which may contribute to the low coancestry of these populations both with each other and with our other populations, as well as to observed patterns of neutral and adaptive phenotypic divergence. Regardless of where the genetic variation underlying trait values originates, the concordant trait and habitat differences between these low elevation populations and the other teosinte populations in excess of neutral expectation drives the strong signal of adaptive trait divergence.

**Figure 4:**
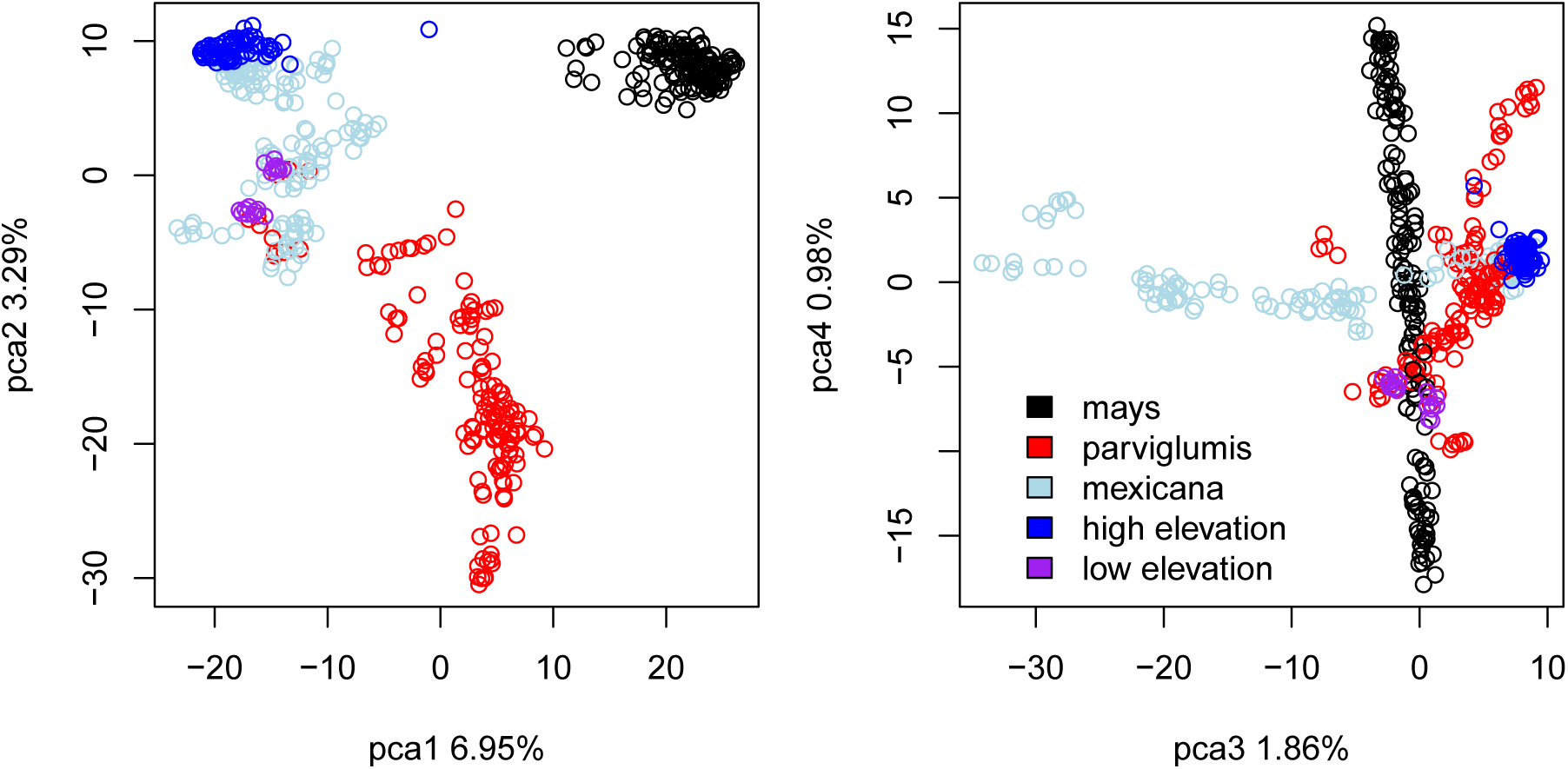
Paired comparisons of the first four axes of the PCA analysis of genotypes comparing individuals (points) in the study populations (purple for low elevation populations and blue for high elevation) to individuals from diverse populations of each subspecies in the genus *Zea*, including individuals from ssp. *mays* (black), ssp. *parviglumis* (red), and ssp. *mexicana* (light blue). Percentages indicate amount of SNP variation explained by that PCA axis.

### Biota alter additive genetic variance and covariance

To test whether rhizosphere biota alter genetic variance and covariance, we fit G matrices to phenotypic data across our 30 population and rhizosphere biota combinations (ten populations in each of three soils). We then used eigentensor analysis (Aguirre et al., 2014) to test for variation among the G matrices and to determine which subspaces of the G matrix are responsible for any variation, and applied selection gradients to asses the biological impact of variation.

We find variation across the G matrix set, largely due to differences in covariance between plant size and phenology. The first 13 of the 15 eigentensors explain significantly more variance than eigentensors calculated on G matrices built from randomly shuffled breeding values (see Figure S4), meaning that there is significant variation in our set of G matrices. Each eigentensor consists of a matrix indicating contributions of individual cells in the G matrix to the eigentensor. The first and second eigentensors explain the largest portions of the variation (43% and 17%, respectively), and reflect primarily covariance between flowering time and each of stem width, root biomass, and shoot biomass, as well as covariance among size traits, (Figure 5).

**Figure 5:**
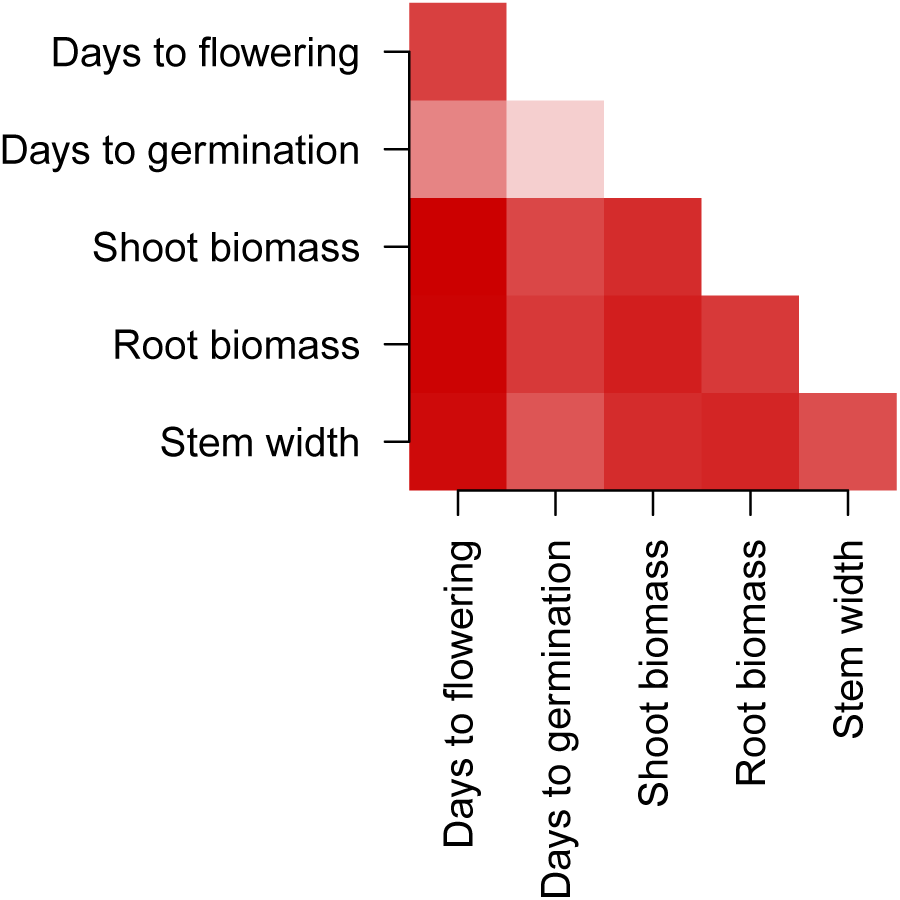
The first eigentensor of the set of G matrices. This eigentensor shows the contribution of each element of the G matrix (symmetric around the diagonal) to the divergence of the set of G matrices. Darker colors indicate greater contributions.

Decomposing the eigentensors into eigenvectors indicates the contribution of each trait to variance and covariance explained by the eigentensor. For both the first and second eigentensor, the majority of the variation explained can be found in the first (84% and 55%) and second (11% and 39%) eigenvectors. One leading eigenvector of the first two eigentensors implicates correlated changes in size and phenology elements of G matrices, whereas phenology and size traits have opposite signs of correlation to the other leading eigenvectors, implicating contrasting changes (Figure S8). Variation among G matrices reflects eigenvector patterns, and includes both differences in estimated strength of variance-covariance and in the sign of covariance between phenology and size traits (Figure 6, and Figure S5).

**Figure 6:**
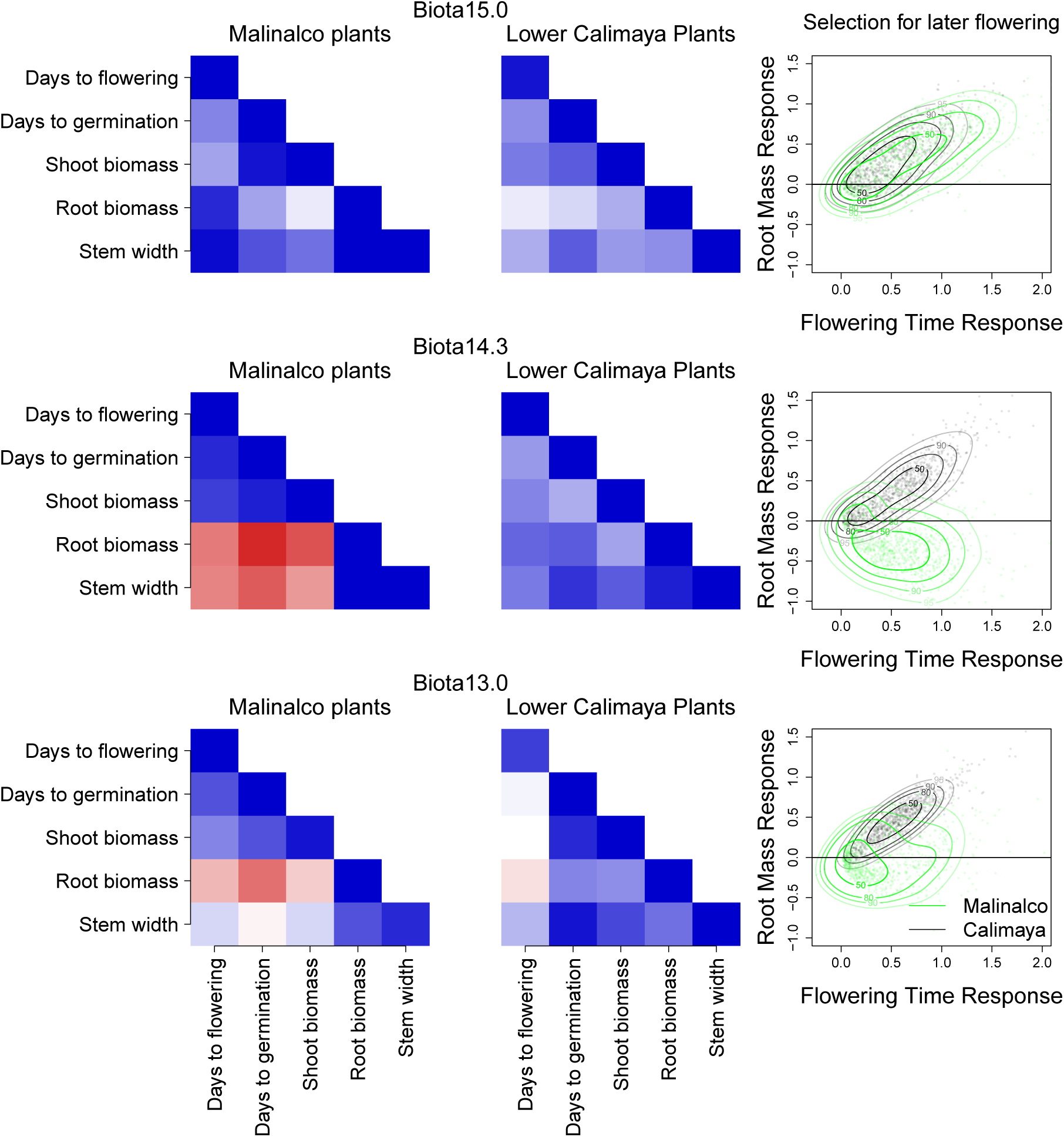
G matrices estimated for populations Malinalco and Lower Calimaya (left columns) across each biota (rows). Redder values indicate more negative genetic covariance, and bluer values indicate more positive additive genetic covariance (off-diagonal) and variance (diagonal). Projected responses of flowering time and root biomass (right column) for G matrices in each biota to selection on flowering time (Malinalco in green, Lower Calimaya in black), with fitted bivariate probability density kernels at 50%, 80%, 90%, and 95% (lines) for responses across MCMC G matrix samples (points).

If differences in additive genetic variance and covariance in traits are large enough, we might predict that populations would respond differently to selection depending on interactions with rhizosphere biota. To predict whether variation in G matrices was biologically meaningful, we simulated responses of teosinte trait means to selection using a selection gradient. Briefly, we projected G matrices onto the same selection gradient vector using the multivariate breeder’s equation (Lande, 1979, 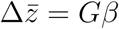) across the posterior distribution for each G matrix. We chose to evaluate selection for later flowering because covariance with flowering time was identified as the greatest axis of variation among the G matrices in our eigentensor analysis. Our selection gradient *β* represents selection for later flowering time of arbitrary strength (equivalent to 1 standard deviation change in pre-reproduction mean phenotype), and without direct selection on any of the other traits in the G matrix. However, responses to selection will reflect the genetic covariance between other traits and flowering time as well as the additive genetic variance for flowering time. We report the predicted response of the trait with the next highest influence on the first eigentensor (root biomass).

The selection gradient approach revealed that size traits were generally predicted to increase with selection for later flowering, but biota altered this in a few cases. Most predicted responses to selection overlapped both across populations and within populations across biota. However, the low elevation population Malinalco was predicted to respond to selection for later flowering quite differently depending on which biota it grew with (both compared to itself in other biota and to other populations in the same biota). Malinalco has an especially divergent matrix in Biota14.3, where it responds with decreases in size traits with selection for later flowering, compared to Biota15.0 where it responds with increases in size traits (Figure 6). One additional population (Texcoco, mid-elevation) lacks a predicted increase in size with later flowering selection only in Biota14.3, and responses of some other populations differ across biota in unique ways (Figure S6, Figure S5). The dramatic differences across biota for Malinalco likely contribute to the significant variation detected by eigentensor analysis.

## Discussion

Abiotic environments can shape evolution of phenotypes across landscapes, such as for flowering time and branching in the sticky cinquefoil across climate and elevation in California (Clausen et al., 1947). Species interactions can also dramatically alter phenotypes: for example, predator presence determines morph development in animals such as the water flea (Tollrian, 1995) and the Trinidadian guppy (Gosline and Rodd, 2008; Ruell et al., 2013), and likewise, shading from competitors shifts morphological development across plants (Smith, 1982; Schmitt, 1997). Yet, we still know little about how biotic interactions may interact with phenotypic adaptation across abiotic environments (but see, Wagner et al., 2014). Here, we found that several teosinte phenotypes were shaped by both environmentally determined divergent selection in teosinte and plastic effects in response to biota. We further determined that shifts in additive genetic variance and covariance due to interactions with rhizosphere biota could alter the course of future phenotypic evolution in teosinte.

### Plastic responses and trait divergence

Plastic trait expression in response to the environment at both the level of the whole organism (Falconer, 1952; West-Eberhard, 1989) and individual genes (Hunter, 2005; Des Marais et al., 2013) are ubiquitous and well-characterized drivers of phenotypic differences across environments. Nonetheless, studies of phenotypic divergence across populations in a wide array of species have revealed that trait variation across populations is often shaped largely by natural selection (Kawecki and Ebert, 2004; Leinonen et al., 2013). In our greenhouse common environment, biota plastically altered the expression of a number of traits, and the direction and extent to which rhizosphere biota altered trait expression depended on the teosinte population (Figure 1, Table 1). However, we find strong evidence of adaptive divergence among teosinte populations, particularly for flowering time and root biomass, regardless of the rhizosphere biota applied.

The forces shaping phenotypic divergence among populations are difficult to distinguish (Karhunen et al., 2014). One way to test for the influence of environment is to compare phenotypic similarity among populations with environmental similarity, all while accounting for the amount of phenotypic similarity we expect among populations simply due to genetic similarity (Karhunen et al., 2014). We find here that environments pattern variation in teosinte phenotypes much more strongly than we expect given genetic similarity among populations, implicating environment-driven selection on teosinte traits. The targets of selection are more likely to be traits for which we find the strongest signals of selection (days to flowering, root biomass, Figure 3), and the agents of selection are more likely to be environmental variables with the strongest correlations to traits (mean annual temperature, soil water holding capacity, Table 1, Figure 1). Yet, it is also possible that these are merely closely correlated to the true targets and agents of selection (Karhunen et al., 2014). Similarly, while we infer that traits with weaker or no signal of environmental selection may not be under selection, these traits may only exhibit divergence in other environments. For example, divergent selection in maize and teosinte has been implicated in stem color variation in other studies (Hufford et al., 2013), and though we find only weak support for divergent selection (see Figure S3), heritable differences in stem color may be expressed more strongly in cold environments (such as the conditions in Hufford et al., 2013).

Gene flow between teosinte subspecies may have contributed to neutral and adaptive trait divergence. Introgression can sometimes underlie trait adaptation to both abiotic gradients and biotic interactions (Whitney et al., 2006, 2010; Song et al., 2011; Pardo-Diaz et al., 2012; Norris et al., 2015), with resulting trait values ranging from intermediate to transgressive (Whitney et al., 2010). Our analysis suggests populations from warmer sites may share some genetic diversity with the lowland subspecies (*Zea mays* ssp. *parviglumis*, Figure 4), and some traits in our lowland populations shift towards ssp. *parviglumis* values.

Our observations of trait divergence themselves varied, with different traits and populations showing evidence of divergence depending on the rhizosphere biota with which plants were inoculated (Figure 3). Several phenomena could explain this interactive effect of plant genotype and rhizosphere biota. Community composition of rhizosphere biota differs strongly across abiotic source conditions and also shifts in response to plant genotype (Bongers and Ferris, 1999; Bulgarelli et al., 2012; Palomares-Rius et al., 2012; Peiffer et al., 2013; Lebeis et al., 2015; Walters et al., 2018; Erlandson et al., 2018), both of which vary across our populations (The authors, 2018, Figure 2). In turn, changes in biota community composition often produce changes in plant phenotypes (Klironomos, 2002; Lau and Lennon, 2012; Wagner et al., 2014). Alternatively, variable effects of biota on plant traits may be a product of selection, especially since rhizosphere bacteria can respond to selection on host phenotypes faster than hosts themselves through changes in community composition (Lau and Lennon, 2012). Moreover, rhizosphere biota have previously been found to alter both flowering time and the fitness consequences of flowering time (Wagner et al., 2014). Consistent with this, we found that biota simultaneously alter flowering time expression (Figure 1, Table 1) and genetic correlations between flowering time and shoot biomass (Figures 6 and S5), which itself is tightly correlated to teosinte fitness (data from Piperno et al., 2015, Figure S10).

### Variation in G matrices - potential responses to changing environments

The course of adaptation to divergent environments can be strongly affected by trait variance and covariance (Schluter, 1996; Etterson and Shaw, 2001; Chenoweth et al., 2010). For example, differences between the direction of selection and the major axis of trait variation may cause trait divergence among populations to reflect the major axis of the G matrix in individual populations (Schluter, 1996; Chenoweth et al., 2010; McGlothlin et al., 2018). However, changing environmental conditions may simultaneously shift both selection on traits (Etterson and Shaw, 2001) and trait variance-covariance relationships (Wood and Brodie III, 2015). The combination of these two processes can lead to unpredictable side effects for trait evolution (Wood and Brodie III, 2016). Studies of G matrix shifts across environments are few because accuracy in G matrix estimates requires large sample sizes (e.g. our ancestral G matrix in the divergence analysis, see also Puentes et al., 2016), and multiplying across populations and environments becomes intractable. Using eigentensor analysis, we leverage many estimates of G matrices from fewer families and individuals. We detect significant variation in additive genetic variance and covariance of size and phenology between teosinte populations and across different biota (Figures S4, 5 & S8), despite low confidence in any one G matrix estimate (see wide variance of posterior estimates Figures S7 and S9).

The differences we observe in G matrices across biota and populations could have arisen due to drift, selection, or genotype-by-biota interactions and cryptic variation. Neutral genetic diversity in teosinte populations differs across its range due to demography (Pyhäjärvi et al., 2013; Aguirre-Liguori et al., 2017), and would be expected to generate differences in total additive genetic variation but not G matrix orientation (Roff, 2000; Puentes et al., 2016), but see (Steppan et al., 2002; Roff et al., 2012). Our populations do not differ substantially in total additive genetic variation (see Figure S9), but rather in orientation of the G matrix, suggesting a role for other processes such as introgression (Guillaume and Whitlock, 2007; Parsons et al., 2011), selection (Arnold et al., 2008), and plasticity (Wood and Brodie III, 2015). Introgression from *parviglumis* should align the axis of greatest trait variation with divergence among the subspecies (Guillaume and Whitlock, 2007). In contrast, selection can, at least temporarily, align it with the multivariate selection gradient (Orr and Betancourt, 2001; Walter et al., 2018), consistent with what we see in teosinte: most populations in most biota have positive average covariance for root mass and flowering time, but for others there is negative or no covariance (Figures 6, S5), and we found that these same two traits were shaped by selection across environments. Differences in estimated G matrices can also arise from plastic responses to the environment in which traits are measured: shifting environments can expose genetic variants that do not have equivalent effects across environments (GxE loci, cryptic variation, see McGuigan and Sgrò, 2009; Des Marais et al., 2013; Paaby and Rockman, 2014). Indeed, most differences in our G matrices depend on biota, suggesting that the G matrix orientation is shaped strongly by responses to biota.

As climate changes, we might expect selection on phenotypes to extend the patterns of divergence we already see across climate (Etterson and Shaw, 2001), here for later flowering time and larger root biomass (Figure 3). Genetic constraints can stymie responses to selection pressures (Etterson and Shaw, 2001; Agrawal and Stinchcombe, 2009), but constraint can vary substantially across populations (Sheth et al., 2016). Most teosinte populations have positive correlations between these two traits across biota, and thus no constraint to such a selection gradient. However, we find that interactions with rhizosphere biota can modify this constraint, generating negative genetic covariance (Figure 6). Indeed, our selection skewers analysis (Lande, 1979) predicted diverging multivariate responses to selection for later flowering time across teosinte populations and biota. More generally, responses to selection in teosinte could be unpredictable if populations disperse to sites with novel rhizosphere biota communities or if local biota change composition (likely already occurring, Castro et al., 2010).

## Conclusions

We have demonstrated the potential for species interactions to be drivers of the expression of adaptive phenotypes, as well as their probable involvement in the past or future responses to environmental selection on traits. Biotic interactions are expected to play major roles in how species respond to climate change. Here we show that in addition to immediate ecological effects on population traits, changing biotic interactions have the potential to influence evolutionary responses — which are projected to be necessary to prevent extinctions (Shaw and Etterson, 2012). These results suggest that as we continue to study the biotic interactions across populations and environments, we must invest as well in understanding the broader consequences of these changes for adaptive evolution.

## Supporting Information

**Figure S1:**
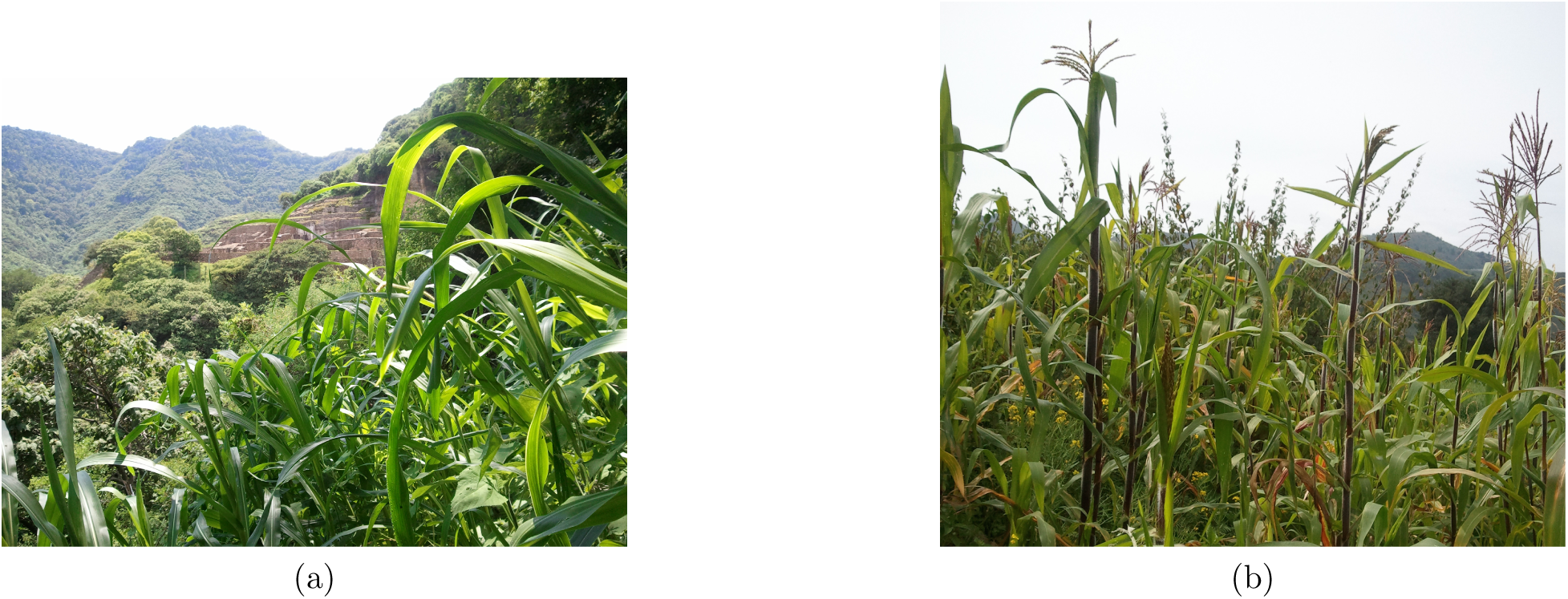
Variation in teosinte stem color in the field. Green at low (a) and red at high (b) elevation. Photographs from the first author.

**Table S1:** Expanded Table 1; best models for traits. The “base” model includes only an intercept with random effects of rhizosphere biota, family, and population. We show the sign and the significance of the slope for simplicity. Second best model variables, slopes and significance are in the last column, except for shoot biomass and stem greenness, where “base” model was best. Abbreviations: TAP for total annual precipitation, MAT for mean annual temperature, SWC for soil water holding capacity (elevation was never in a best or second best model), F family in biota random effect, P population in biota random effect. Symbols: ** pMCMC < 0.05, * pMCMC < 0.1.

| Trait | Main | Random | Slope | DIC base | DIC 1st | DIC 2nd | Effects in 2nd |
| --- | --- | --- | --- | --- | --- | --- | --- |
| Days to flowering | – | F |  | 1713.4 | 1713.0 | 1713.1 | MAT, F, + * * |
| Days to germination | SWC | F+P | +** | 2157.8 | 2131.6 | 2131.6 | TAP, F+P, + |
| Tassel length | SWC | – | +** | 2096.8 | 2096.6 | 2096.6 | SWC, P, + * * |
| Shoot biomass | – | – |  | 2233.3 |  |  |  |
| Root biomass | – | F + P |  | 1889.6 | 1888.7 | 1888.7 | TAP, F+P, + * * |
| Height | MAT | – | – | 2344.7 | 2344.1 | 2344.1 | MAT, F, – |
| Stem width | MAT | P | +** | 2310.0 | 2309.1 | 2309.4 | MAT, F+P, + * * |
| Leaf width | – | F + P |  | 2083.9 | 2083.4 | 2083.5 | TAP, F+P, + |
| Stem greenness % | – | – |  | 2261.6 |  |  |  |

**Figure S2:**
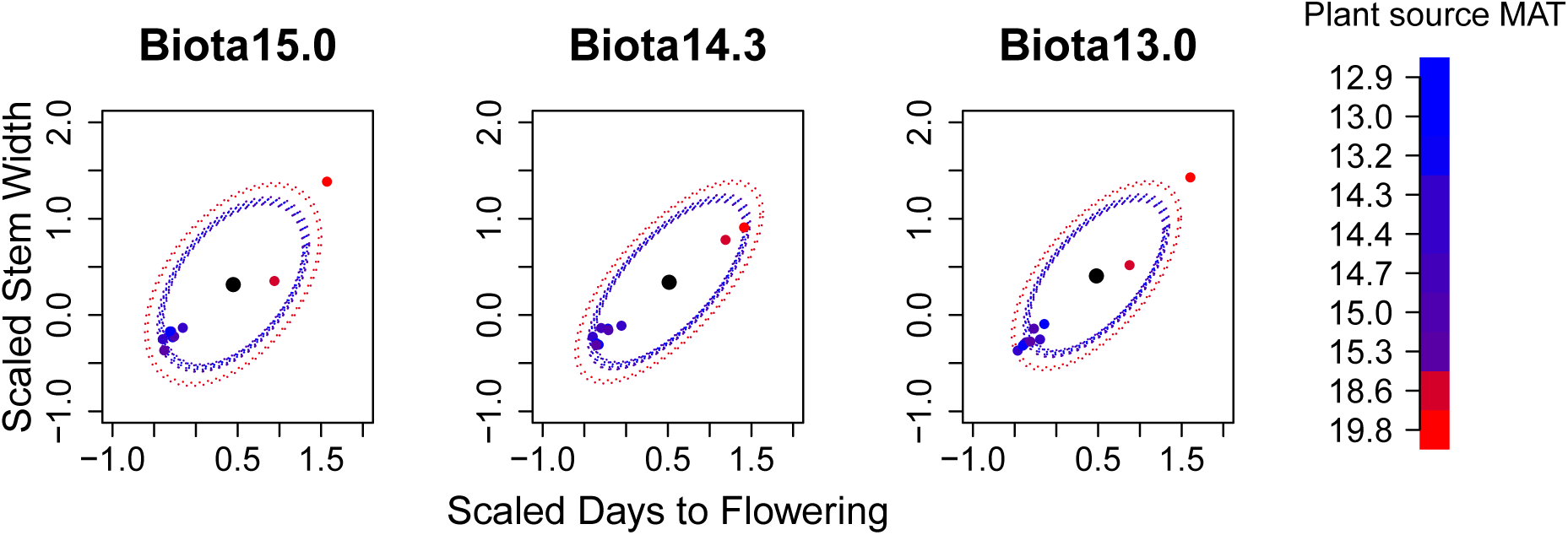
Standardized population means (colored points) and 95% confidence intervals for the neutral expectation of population means (matching colored circles), for flowering day and stem width, estimated by Driftsel. Redder color indicates warmer mean annual temperature at the source site. Means beyond matching confidence intervals indicates significant divergence.

**Figure S3:**
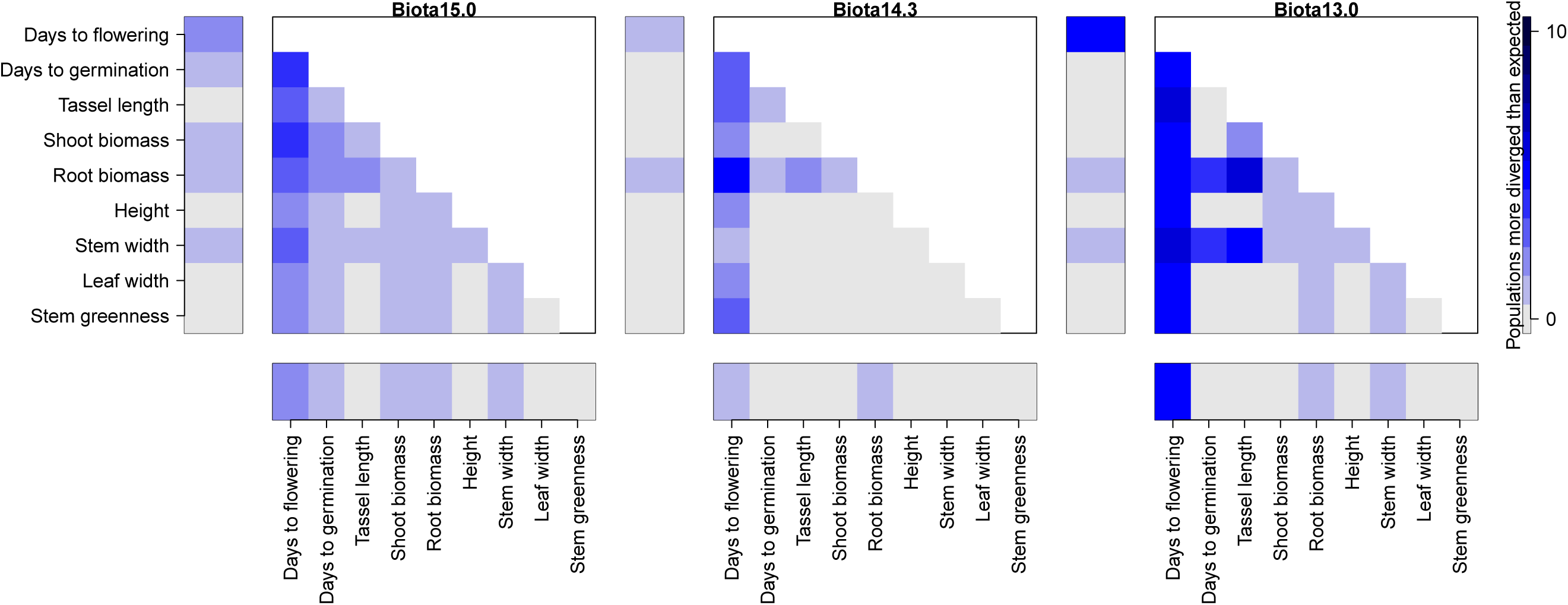
Populations exceeding bivariate expectations of divergence in all biota (separate plots). Darker blue squares indicate trait combinations with more populations outside expectations (each panel of figure S2 is reduced to one square here). Bars on the axes indicate the number of populations in which traits exceed divergence expectations in the univariate case (see Results for higher dimension summary S test).

**Figure S4:**
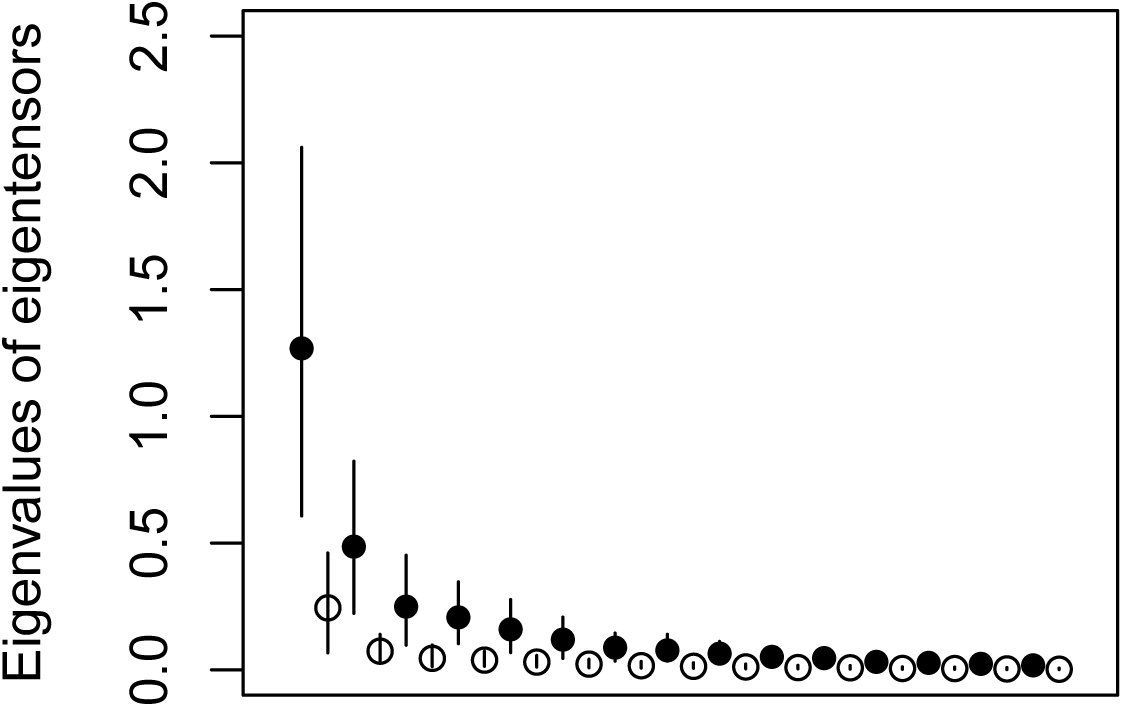
Eigenvalues of the first 11 eigentensors of the set of 30 G matrices (filled points), and the first 11 eigentensors from the randomized array (open points). Error bars represent confidence intervals across MCMC estimations (real set) or across the randomized array.

**Figure S5:**
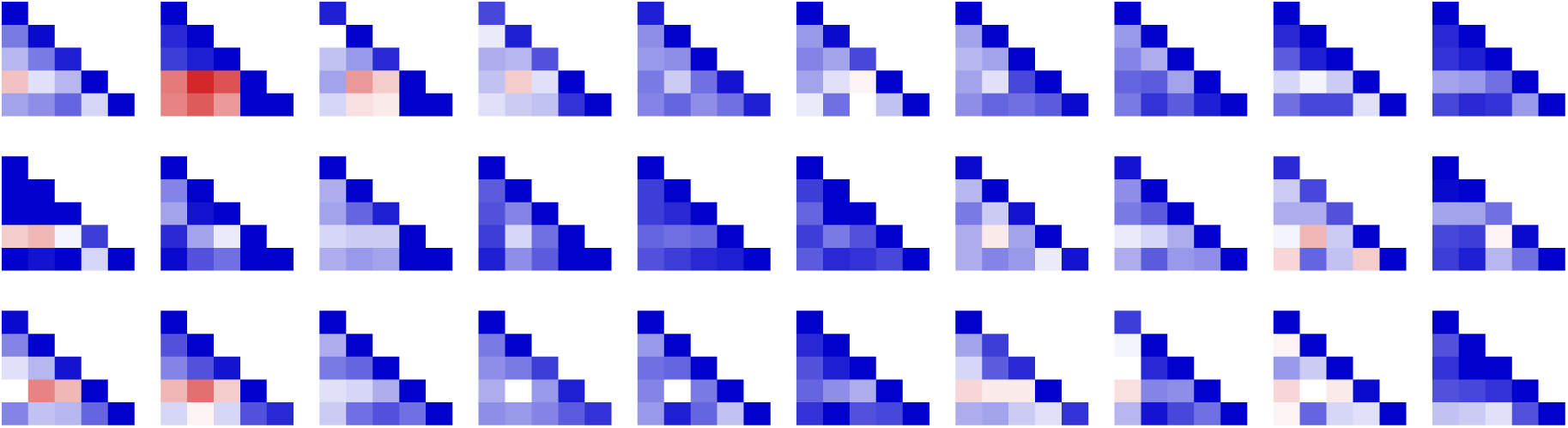
The full set of G matrices across populations (columns) and biota treatments (rows). Colors and trait organization are as in Figure 6.

**Figure S6:**
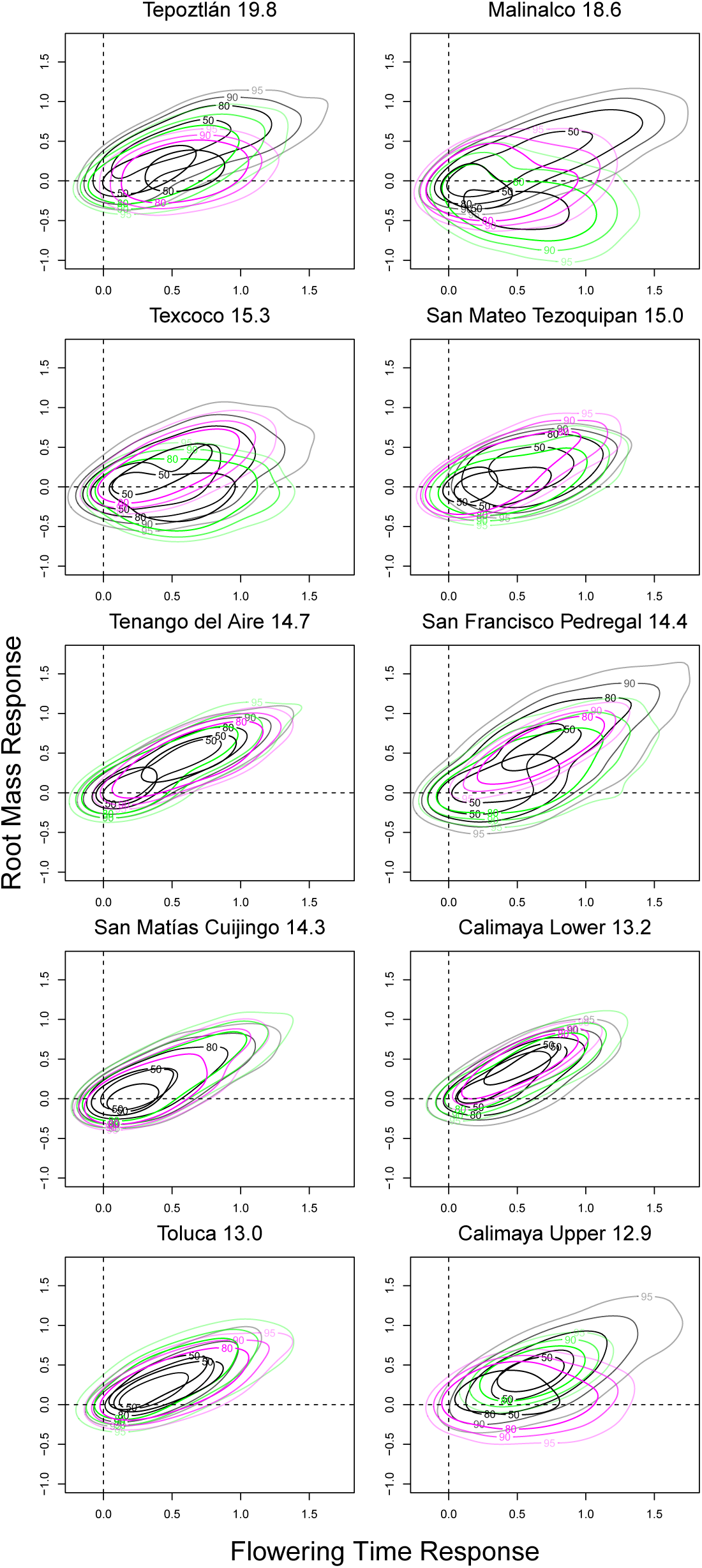
Responses to selection on flowering time for G matrices estimated in each population in different biota. Confidence intervals in contour lines as in Figure 6 (black, Biota15.0; green, Biota14.3; purple, Biota13.0). Dashed lines highlight 0 response for both traits. Plant populations are sorted by MAT °C with names.

**Figure S7:**
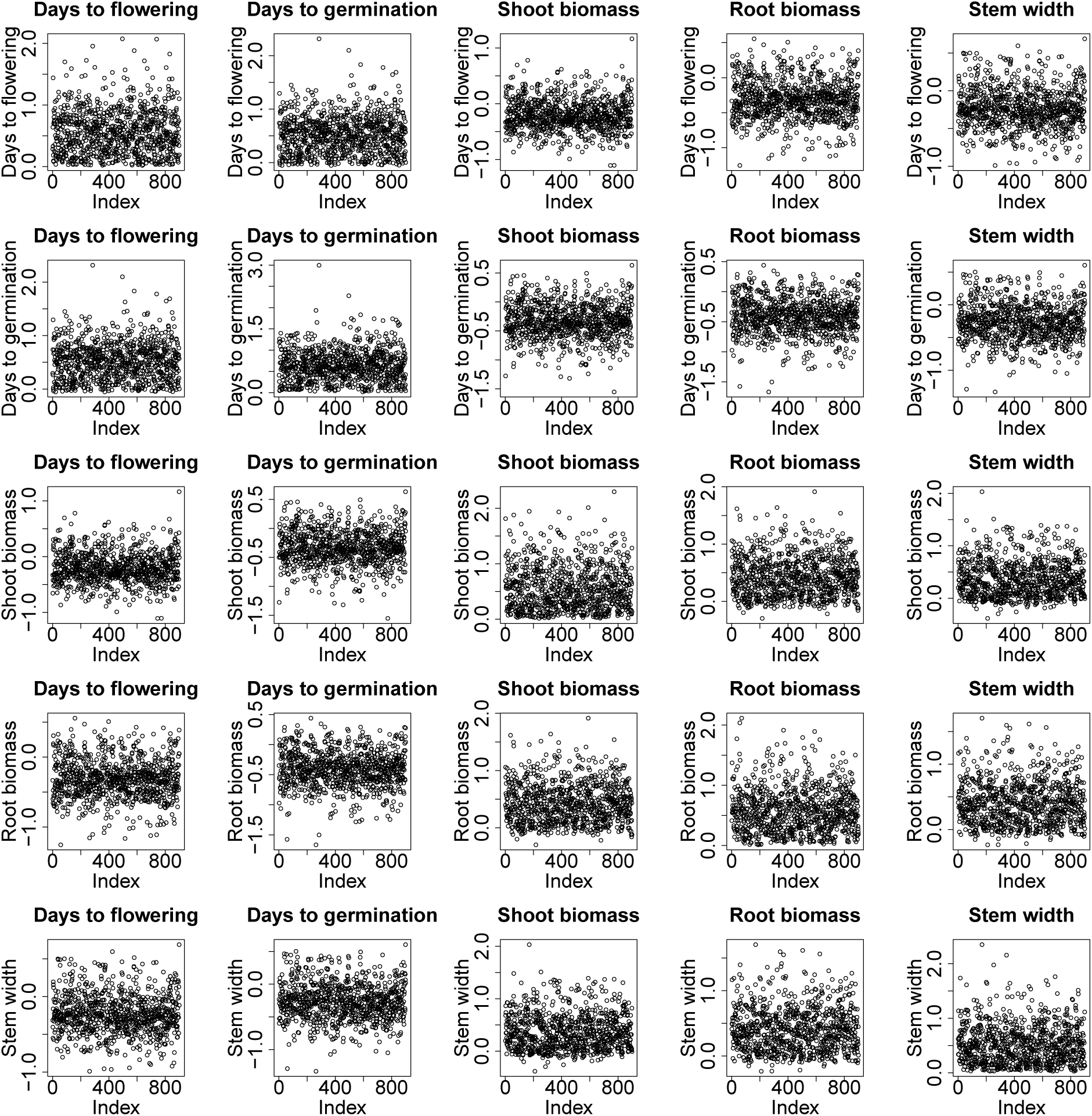
Trace of posterior estimates of the G matrix for Malinalco population in Biota15.0, as an example of MCMC chains.

**Figure S8:**
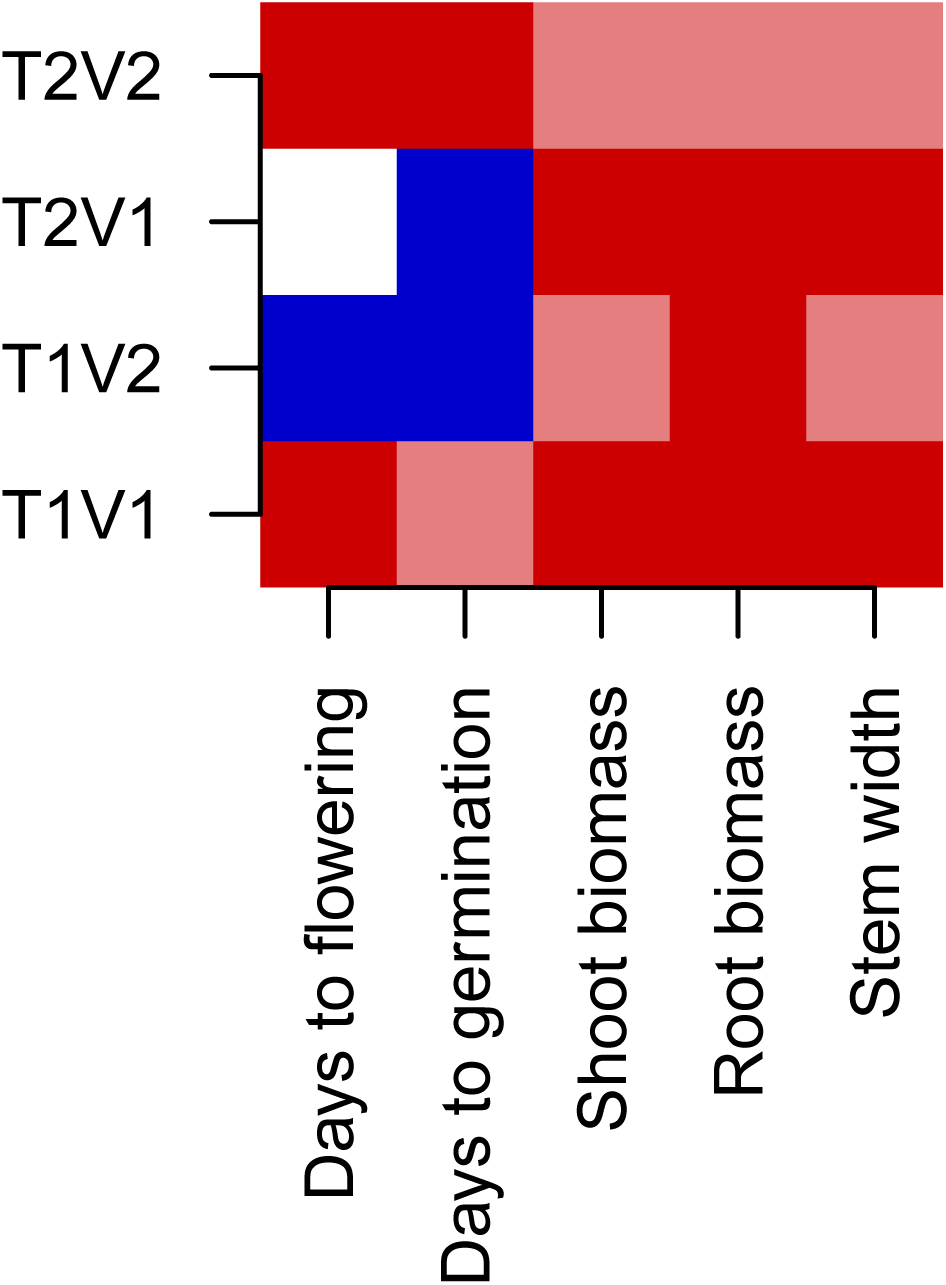
First and second eigenvectors (V1,V2) of the first two eigentensors (T1,T2) of the set of genetic variance-covariance matrices. Red indicates positive loading on the tensor, blue indicates negative loading, and color intensity indicates the strength of loading.

**Figure S9:**
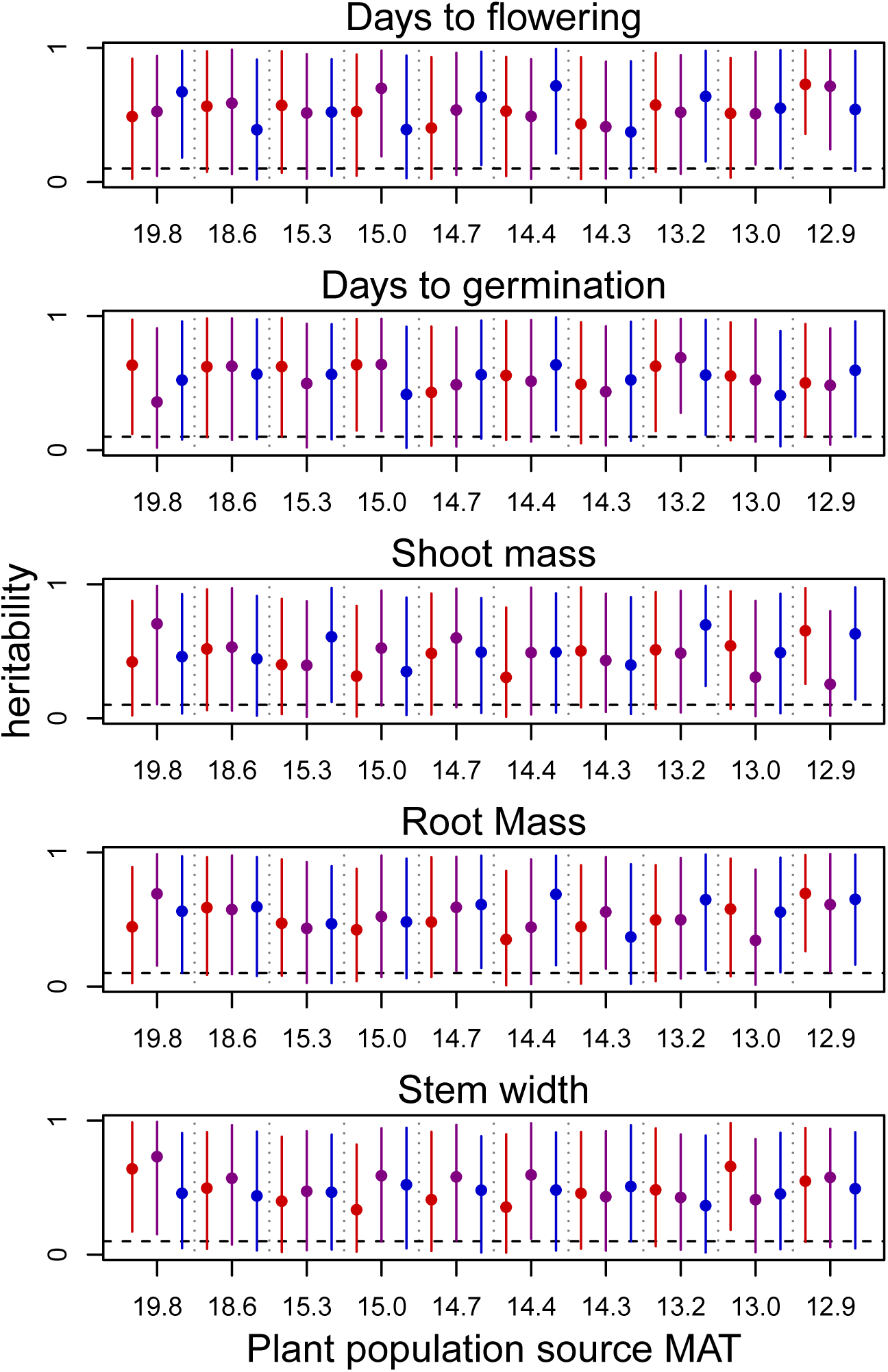
Estimated heritability of each trait in each population calculated from additive genetic variance and residual error (environmental variance) in fitted MCMCglmm models. Points and 95% highest posterior density intervals are colored by the rhizosphere biota in which the plants were measured (red, Biota15.0; purple, Biota14.3; blue, Biota13.0). Vertical dotted lines separate plant populations. Horizontal line is at 0.1 (MCMCglmm estimation cannot produce a value of exactly 0).

**Figure S10:**
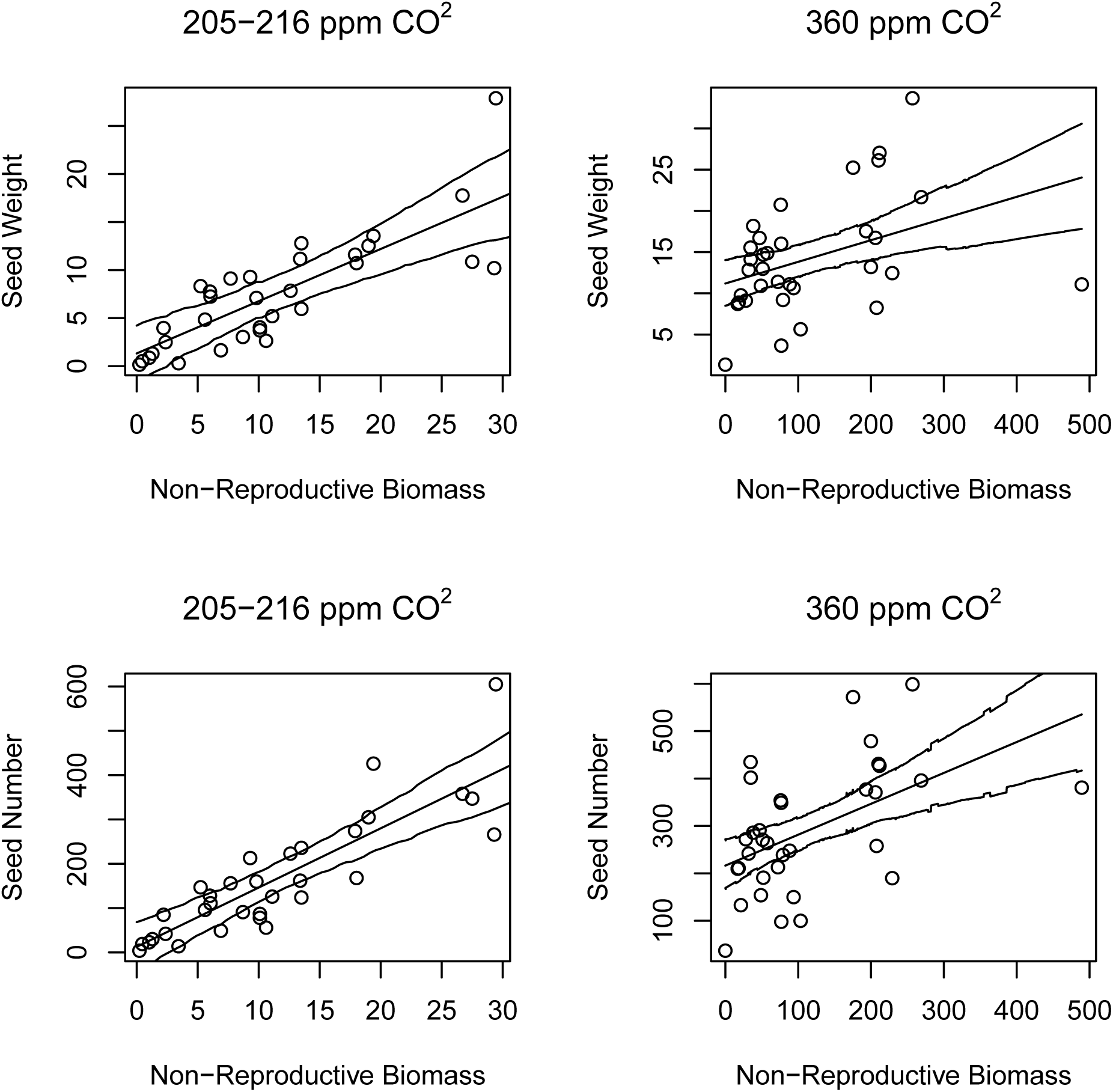
Using raw data reported in Piperno et. al 2015, and provided by Dolores Piperno, we asked if vegetative biomass at flowering (excluding seeds), is correlated to seed mass and number. We used linear models fit with MCMCglmm (as in the main text), including parameters for seed source population and year random effects, as well as for fixed and interaction effects for the CO_2_ treatments. Both seed mass and number were significantly positively correlated to non-seed biomass. (*Y ~ α* + *αyear* + *αpopulation* + *βppm*+*βbiomass*+*βbiomass ppm*) Model predictions are plotted as mean lines surrounded by 95% HPDI intervals. Data is plotted as points. All fitted parameters are significant at pMCMC <0.001

**Author contributions:** All authors contributed substantially to the design of the study, provisioning of materials, and revising of the manuscript. AO proposed the study, collected the data, performed analyses and provided the first draft of the manuscript.
Submitted as an original article to Evolution

## Acknowledgements

The authors would like to thank Jaime Gasca Pineda & Luis Eguiarte and the Eguiarte laboratory for help with field collections, Aida Odette Avendaño-Vázquez, Carlos Fabián de la Cruz, Abenamar Gordillo Hidalgo, Dario Alvarez & Arturo Chavez for assistance in the greenhouse. We thank J. Schmitt and M. Frederickson for comments on the manuscript. The project was funded by UC MEXUS, the UC Davis Center for Population Biology grants to AO, USDA Hatch project CA-D-PLS-2066-H to JRI, and NSF grant IOS-0922703. AO was supported by the NSF GRFP grant DGE-1148897, NSF grant DEB-0919559, and a GSR fellowship from the UC Davis Department of Plant Sciences.

